# Accelerated Bone Healing in Calvarial and Femoral Defects with Injectable Microcarriers that Mimic the Osteogenic Niche

**DOI:** 10.1101/2021.11.05.467478

**Authors:** Candice Haase, Sravani Jaligama, Eli Mondragon, Simin Pan, Eoin P. McNeill, Cynthia Co, Daniel Tahan, Bret H. Clough, Nick Sears, Abhishai Dominic, Jun Kameoka, Carl A. Gregory, Roland Kaunas

## Abstract

Osteo-enhanced human mesenchymal stem cells (OEhMSCs) secrete an osteogenic cell matrix (OCM) that mimics the composition of anabolic bone tissue and strongly enhances OEhMSC retention and subsequent bone repair *in vivo*. Here we demonstrate a system for rapid production of gelatin methacrylate microcarriers coated with decellularized OCM (OCM-GelMA) to serve as an injectable bone graft material with high osteogenic potential comparable to a clinically utilized gold standard, bone morphogenic protein 2 (BMP-2). OEhMSCs seeded onto OCM-GelMA secreted high levels of osteogenic and angiogenic cytokines and expressed higher levels of BMP-2 relative to OEhMSCs on bare GelMA microcarriers. OEhMSCs co-administered with OCM-GelMA microcarriers resulted in enhanced healing of murine critical-sized calvarial defects, which was comparable to that achieved with a BMP-2-laden gelatin sponge control. When tested in a murine femoral defect model, OCM-GelMA co-administered with OEhMSCs also induced profound bone growth within the defect. We submit that OCM-GelMA promotes OEhMSC paracrine release to accelerate bone repair, indicating their potential as a bone graft for use in minimally invasive surgery.

## Introduction

The extracellular matrix (ECM) in tissues serves as both a mechanical support for adherent cells and a repository of bioactive cues that regulate cell function. Synthetic scaffolds have been designed to mimic many biochemical and biophysical properties of native ECM, but are inherently less complex than natural ECM [1]. Consequently, there has been substantial interest in creating biological scaffolds composed of ECM in the form of decellularized tissues. In the case of bone, decellularized bone matrix (DBM) obtained from cadavers has been widely used due to its three-dimensional (3D) structure, osteoinductivity, and similarity to native bone ECM [2]. Although DBM has superior bone regenerative properties to some commonly used bone graft materials, such as tricalcium phosphate and decellularized xenogenic bone, it does not match the efficacy of autologous bone [3]. Autologous bone has its drawbacks as well, including donor site morbidity, secondary surgery, and infection [4].

Decellularized ECM derived from cultured cells offers an alternative to autograft and DBM for creating bone grafts. Decellularized ECM with osteogenic properties has been deposited onto various scaffold materials [5]. The composition of the cell-derived matrix is dependent on cell type and, in the case of hMSCs, their differentiation status. For example, hMSCs treated with a PPAR-gamma inhibitor (GW9662) to promote osteogenic differentiation into OEhMSCs and generation of an osteogenic ECM enriched in collagens VI and XII [6, 7]. Collagens VI and XII are also upregulated in regenerating bone and possess potent osteogenic properties [8, 9]. Importantly, ECM generated by OEhMSCs strongly accelerates bone healing in rodent calvarial [6, 7] and femoral [10] defects, as well as spine fusion models [11].

Methods designed for obtaining grafts containing cell-derived ECM include mechanical harvesting ECM from monolayer culture [7] and deposition of ECM onto implantable scaffolds such as polymer mesh [12], 3D printed scaffolds [13], titanium mesh [14], gelatin sponge [10] and hydroxyapatite microparticles [15]. While each of these studies have demonstrated bone formation, the methods used to produce the ECM are most appropriate for small-scale production with manual culture techniques. More efficient strategies will be required for cell-derived ECM to be a practical strategy for fabricating bone grafts at clinically relevant quantities in a reliable and cost-efficient manner.

A well-established strategy to expand hMSCs in large numbers involves suspension culture on commercial microcarriers using wave-mixed and stirred tank bioreactors [16]. Commercial microcarriers are cytocompatible but are not designed for use as an implantable biomaterial. Gelatin methacrylate is a promising biomaterial for use in tissue engineering due to its biocompatibility, ease of fabrication and ability to support cell attachment and growth [17]. Planar microfluidics are typically used to fabricate GelMA microcarriers and grow cells in static culture for applications such as cardiac cell [18] and hMSC delivery [19] due to their relative simplicity of construction. Throughput is constrained in planar microfluidic devices due to destabilization of droplet formation at high flow rates and delamination of assembled layers of the device at high pressures [20]. Planar devices produce monodisperse droplets only when operated at low values of Reynolds and capillary numbers, while 3D axi-symmetric systems are capable of forming monodisperse droplets at high values of Reynolds and Weber numbers [21].

In the present study, we developed a strategy for facile production of monodisperse GelMA microcarriers using a 3D axi-symmetric microfluidic device and subsequent deposition of osteogenic OCM from hMSCs treated with GW9662 in suspension culture. Following decellularization, fresh hMSCs were cultured on the OCM-coated microcarriers to test the hypothesis that coating the GelMA microcarriers with OCM promotes a pro-regenerative phenotype. Further, we tested the hypothesis that OCM-GelMA microcarriers accelerate the formation of mature bone in both calvarial and long bone critical size defects.

## MATERIAL AND METHODS

### Device Fabrication and Assembly

The 3D flow-focusing device was fabricated from HTM140 M resin (EnvisionTEC) using a high-resolution stereolithographic 3D printer (EnvisionTEC) as previously described with slight variation [35]. The internal dimensions were modified with the orifice being 250 µm in diameter.

### GelMA Synthesis

Methacrylation of gelatin was performed as previously reported to obtain GelMA with 80% methacrylation [17]. Briefly, type A porcine skin gelatin (Sigma-Aldrich) was mixed at 10% (w/v) into Dulbecco’s phosphate buffered saline (DPBS; GIBCO) at 60°C and stirred until fully dissolved. Methacrylic anhydride was added to the gelatin solution until reaching the target volume at a rate of 0.2 mL/min while stirring at 60 °C and then allowed to react for 1 h. Following a 5X dilution with additional warm (40 °C) DPBS to stop the reaction, the mixture was dialyzed against distilled water using dialysis tubing (10 kDa cutoff) for 1 week at 60 °C to remove salts and unreacted methacrylic acid. The solution was lyophilized for 1 week to generate a white porous foam and stored at -80 °C until further use.

### Microcarrier Fabrication

An aqueous solution of 7.5%(w/v) GelMA and 1%(w/v) lithium phenyl-2,4,6-trimethylbenzoylphosphinate in PBS at 50°C and an oil solution of mineral oil solution containing 0.25%(v/v) concentration of Span 80 (Sigma Aldrich) converge in the flow-focusing device (Figure 1A) with flows driven by two syringe pumps. At the junction of these two coaxial flows, fluid shear forces from the mineral oil flow promote periodic separation of discrete droplets. To initiate polymerization, the droplets repeatedly flowed past a Gen 2 Emitter (LED Engin, San Jose, CA) for a total of 50 s of exposure to 75 mW/cm^2^ UV light at 365nm. All the previous steps were performed within an insulated enclosure maintained at 50°C. The polymerized microcarriers were then collected in a cold PBS bath, washed three times at 3000 rpm for 10 min to remove oil, and stored in PBS at 4°C.

**Figure 1.**
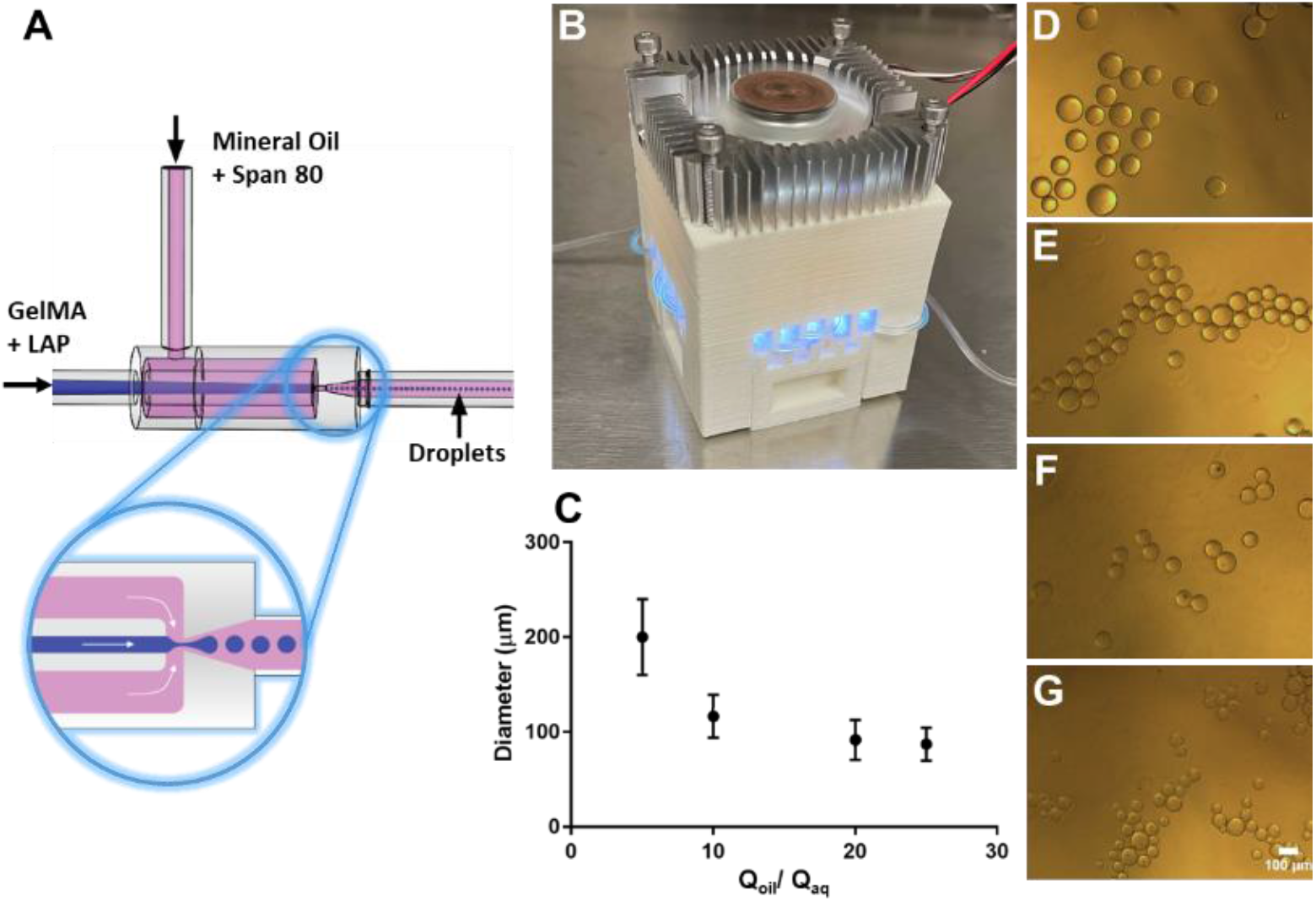
Generation of photocrosslinked GelMA microcarriers. A) Schematic representation of the 3D printed flow-focusing device used to generate GelMA droplets within a mineral oil carrier solution. B) Custom UV light chamber for photocrosslinking. C) Plot summarizing the relationship between droplet diameter and *Q*_oil_/*Q*_aq_ relationship with representative bright-field images of microcarriers produced with *Q*_oil_/*Q*_aq_ ratios (mean±SD) of 5 (D), 10 (E), 20 (F) and 25 (G). Scale bar is 100 μm.

### SEM Imaging

A scanning electron microscope (SEM, Tescan Vega 3) was used to inspect the surface morphology of the bare and OCM-coated microcarriers. The microcarriers in PBS were injected into a polycarbonate filter holder (Cole-Parmer) containing a Grade 415 filter paper (VWR) to remove excess PBS. The microcarriers were then dehydrated using ethanol/water mixtures of 5% (v/v) increments up to 100%. The microcarriers were washed three times with 100% ethanol to remove remaining water. Then, hexamethyldisilazane/ethanol mixtures (HMDS, Sigma-Aldrich) of 5% increments up to 100% were used to gradually replace the ethanol in the microcarriers with HMDS followed by three washes with 100% HMDS. The microcarriers were left to dry in a chemical hood for 96 hrs to slowly evaporate the HMDS without collapsing the microcarriers.

### hMSC culture

hMSCs were acquired from Texas A&M Health Science Center adult stem cell distribution facility in accordance with institutionally approved protocols. Prior to culture on microcarriers, hMSCs were cultured according to standard protocols [7]. All reagents were purchased from Invitrogen Corporation (Carlsbad, CA) unless otherwise stated. Briefly, hMSCs were thawed from a cryopreserved stock and plated on standard tissue culture plastic at approximately 5,000 cells per cm^2^, in complete culture medium (CCM) consisting of alpha minimum essential media, 20% fetal bovine serum (Atlanta Biologicals (Norcross GA), 100 units/ml penicillin, and 100 µg/ml streptomycin, and 4 mM of additional glutamine supplementation. After 24 hr in a humidified incubator at 37°C in 5% (v/v) CO2, cells were then lifted by trypsinization and plated at 500 cells per cm2. Cells were cultured with change of media every 2 days until 60-70% confluency (10,000-15,000 cells per cm^2^) was attained. This was designated one passage of hMSC expansion. For generation of OEhMSCs, media was exchanging to Osteogenic Base Media (OBM, CCM containing 5mM β-glycerophosphate and 50 µg/ml ascorbic acid) supplemented with 10uM GW9662 and the media was exchanged every 2 days thereafter over a total of 10 days. Passage 4 or 5 hMSCs were used in *in vitro* experiments, while passage 2 OEhMSCs were used in animal studies.

### OCM-coating of microcarriers

hMSCs were combined with microcarriers in 10ml of CCM in low-adherence 6-well plates and allowed to adhere to the microcarriers for 2 hr with orbital shaking at 20 rpm. To perform rotating suspension culture, cell-laden microcarriers were transferred to 10-ml Rotating Wall Vessels (RWVs) mounted on a rotation controller (Synthecon model RCCS-8DQ)). After 2 days, media was changed to OBM with 10uM GW9662 to induce deposition of OCM, with the media was exchanged every 2 days thereafter over a total of 10 days. The OCM-coated microcarriers were then processed as previously described for OCM-coated bioinks [22]. Briefly, the OCM-coated microcarriers were washed in excess PBS and decellularized in a lysis buffer (PBS containing 0.1% (v/v) Triton X-100, 1mM MgCl_2_ and 1U/ml DNAse I) for 8 hr at 37°C with orbital shaking at 60 rpm. Decellularized microcarriers were washed sequentially in dH_2_O, acetone, and dH_2_O and then stored in PBS until further use.

### Immunohistochemistry

GelMA and OCM-GelMA microcarriers were each blocked with 5% goat serum (MP Biomedicals) and 0.3% Triton X-100 (Sigma Aldrich) for 1 hr at 25°C. Microcarriers were incubated overnight at 4°C in blocking buffer with rabbit anti-human type VI collagen and rabbit anti-human type XII collagen primary antibodies (each 1:200, Novus Biologicals, Littleton, CO). Microcarriers were washed with PBS and incubated in goat anti-rabbit Fluorescein-conjugated secondary antibody (1:500, Millipore) for 2 hours at 25°C. Finally, samples were washed in PBS and imaged on an upright confocal microscope (Nikon C1 LSC confocal head, Nikon FN1 upright microscope, 20× water-dipping objective, 40-mW Argon ion laser (Melles Griot)). Image stacks were used to produce 3D maximum-intensity projections.

### Live/dead assay

hMSCs were cultured on GelMA and OCM-GelMA microcarriers for 48 hr in CCM, then harvested to assess cell viability. Samples were treated with PBS containing 0.1% BSA (Sigma Aldrich), 1uM calcein AM (AnaSpec), and 7.5uM propidium iodide (Sigma Aldrich) for 45 min at 37°C. The microcarriers were then washed with PBS and stacks of images were collected to produce maximum intensity projections.

### ELISA

Commercially available duoset assay kits for osteoprotegerin (OPG), platelet-derived growth factor-AA (PDGF-AA), vascular endothelial growth factor (VEGF), fibroblast growth factor 2 (FGF-2), dickkopf-related protein-1 (Dkk-1) and transforming growth factor beta 1 (TGF-β1) were obtained from R&D systems (Minneapolis, MN) and performed on conditioned media from hMSCs grown on GelMA and OCM-GelMA microcarriers in RWV bioreactors.

### qRT-PCR

The number of hMSCs present on microcarrier cultures was measured using qRT-PCR for glyceraldehyde-3-phosphate dehydrogenase (GAPDH). Cells were enumerated by comparison with known hMSC standards. Microcarriers were subjected to total RNA extraction using a commercially available kit (High Pure kit, Roche, Indianapolis, IN). Total RNA was then used to make complementary DNA (cDNA) (Superscript III cDNA kit, Invitrogen). Then 0.5 µg of cDNA was amplified in a 20µl reaction containing SYBR Green PCR master mix (Fast SYBR Green, Applied Biosystems) on an AriaMx Real-Time PCR System (Agilent Technologies). Collagen and BMP-2 mRNA expression data was calculated with the 2^-ΔΔ^CT method using human GAPDH as a reference. The PCR primers are described in Table 1.

**Table 1.**
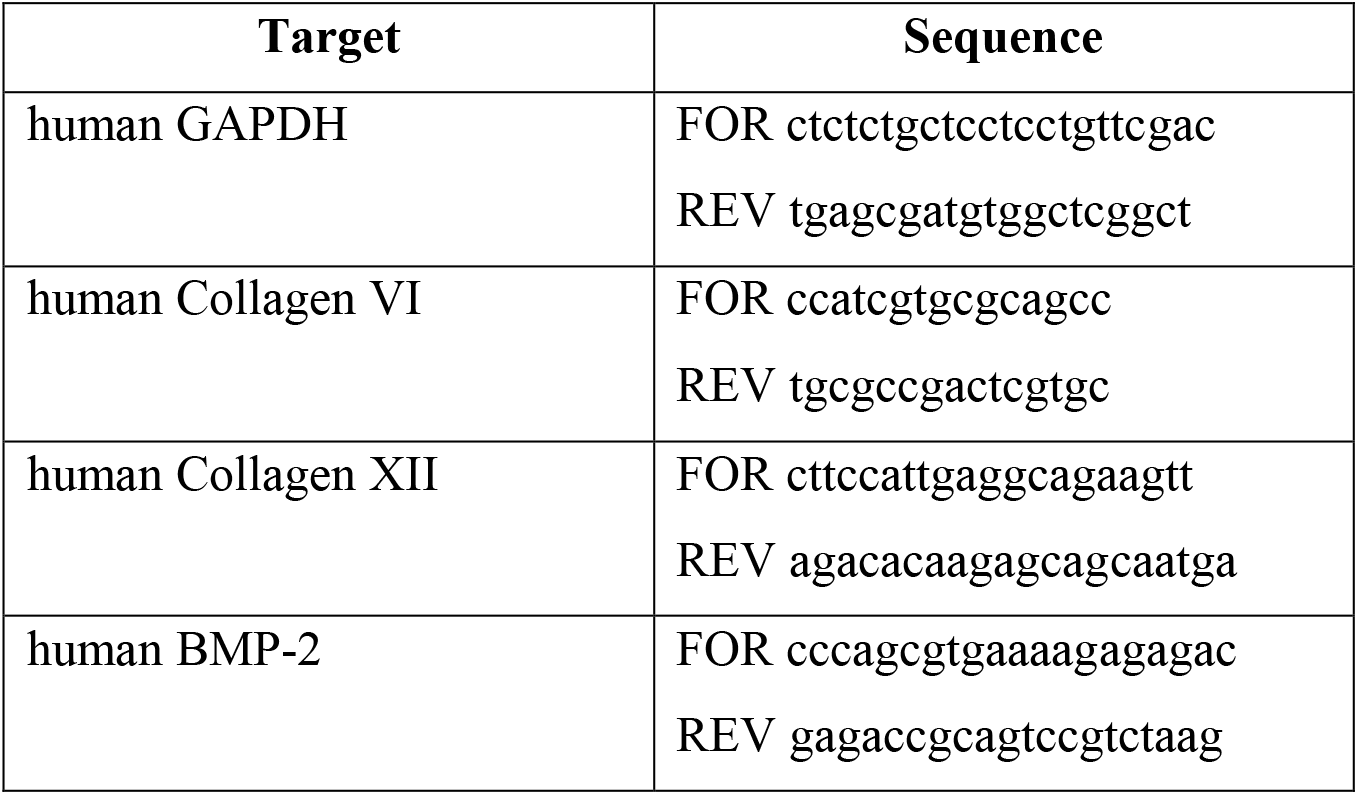
Primer sequences for qRT-PCR.

### Calvarial defect model

Studies were performed in accordance with an institutionally approved animal use protocol as described previously [6, 7]. Briefly, a 4 mm diameter circular full-thickness calvarial defect was generated in immune-compromised nude mice, followed by administration of 50 µL (settled volume) of microcarriers, with or without 2 million OEhMSCs, suspended in a mixture of 25 µL thromboplastin and 25 µL reconstituted human plasma. The plasma was allowed to clot in situ, the scalp was closed by suture and defects healed for 4 weeks. As a positive control, gelatin foam (Gelfoam, Baxter International, Deerfield, IL) saturated with 50 µg BMP-2 (Infuse, Medtronic, Minneapolis, MN) was administered to fill the defect. Mice were then humanely euthanized and calvarial bones were carefully dissected with a dental cutting wheel. Digital x-rays were collected of calvaria and analyzed for densitometry as described previously [7]. A relative healing index (RHI) was calculated, where 1 represented radioopacity equivalent to the contralateral side and 0 is equivalent to air. These calvarial specimens were then fixed and prepared for histology.

### Femoral defect model

Studies were performed in accordance with an institutionally approved animal use protocol. A 3 mm long pin-stabilized defect was generated in 2 month-old immunodeficient nude mice as described previously [10, 23] in accordance with institutionally approved protocols. 25 µL (settled volume) of microcarriers, with or without 1.8 million OEhMSCs, suspended in a mixture of 20 µL of thromboplastin and 20 µL reconstituted human plasma. The entire mixture was administered to the defect and the muscle and skin layers were closed by suture. After 5 weeks of healing, mice were humanely euthanized and hind-limbs were fixed in buffered formalin. Femurs were scanned using a Skyscan1275 micro-computed tomography (µCT) scanner (Bruker, Belgium) as described [23]. Because significant bone formation was observed at the site of the defect and also at distal sites, measurements of bone volume and surface:volume ratios were calculated for the entire femora using CTAnal software (Bruker).

### Histology

Standard histology was performed on decalcified, paraffin-embedded specimens as described previously [7, 10]. Calvarial sections were made in the sagittal plane and femoral sections were made axially after removal of the pin.

### Statistical Analysis

Statistical tests and data plotting were performed with GraphPad Prism version 7.00 for Windows. Specifically, two-tailed t-test was used to determine statistical significance, assuming unequal sample variance for each experimental group when comparing individual groups. For multiple tests of means within data sets, data was statistically analyzed by one-way analysis of variance (ANOVA) and post-analyzed by Tukey’s method.

## RESULTS

### GelMA microcarrier production

The axisymmetric flow-focusing device rapidly produces aqueous droplets of GelMA and LAP photoinitiator carried in mineral oil containing surfactant (Fig. 1A) that then flowed through a section of tubing exposed to ultraviolet irradiation to induce photocrosslinking (Fig. 1B). Increasing the oil-to-aqueous flow rate ratio (Q_oil_/Q_aq_) from 5 to 25 resulted in a decrease in the mean and variance of the droplet diameter (Fig. 1C-G). A flow rate ratio of 25 was used to produce the microcarriers used in subsequent experiments since this resulted in a relatively high rate of production of microcarriers (3×10^4^ min^-1^) with a diameter (87 ± 17μm) small enough to easily passed through an 8 gauge delivery cannula (3.4 mm inner diameter).

### Generation of OCM-Coated GelMA Microcarriers

The overall strategy for generating OCM-coated GelMA (OCM-GelMA) microcarriers is illustrated in Figure 2A. First, crosslinked microcarriers were seeded with hMSCs in low-adhesion plates on an orbital shaker for 2 hr in CCM to promote initial adhesion. The cell-seeded microcarriers were then loaded into rotating wall vessel (RWV) bioreactors for 2 days of expansion in CCM followed by 8 days of osteogenic culture in OBM containing GW9662 during which time the cells deposited OCM. Microcarriers harvested from the RWVs were then decellularized using our previously published methods [22] and imaged to characterize the resulting deposition of OCM. OCM-GelMA microcarriers contained high amounts of fibrillar extracellular matrix (Fig. 2C’, asterisks) and mineralized nodules (Figure 2C’, arrow), whereas the surfaces of GelMA microcarriers were smooth (Figure 2B). We have previously demonstrated that hMSCs monolayers cultured in the presence of GW9662 deposit OCM rich in collagen types VI and XII [7], thus representing useful biomarkers for OCM. Immunostaining showed that collagens VI (Fig. 2D vs 2E) and XII (Fig. 2F vs 2G) were present on the OCM-GelMA, but not GelMA, microcarriers.

**Figure 2.**
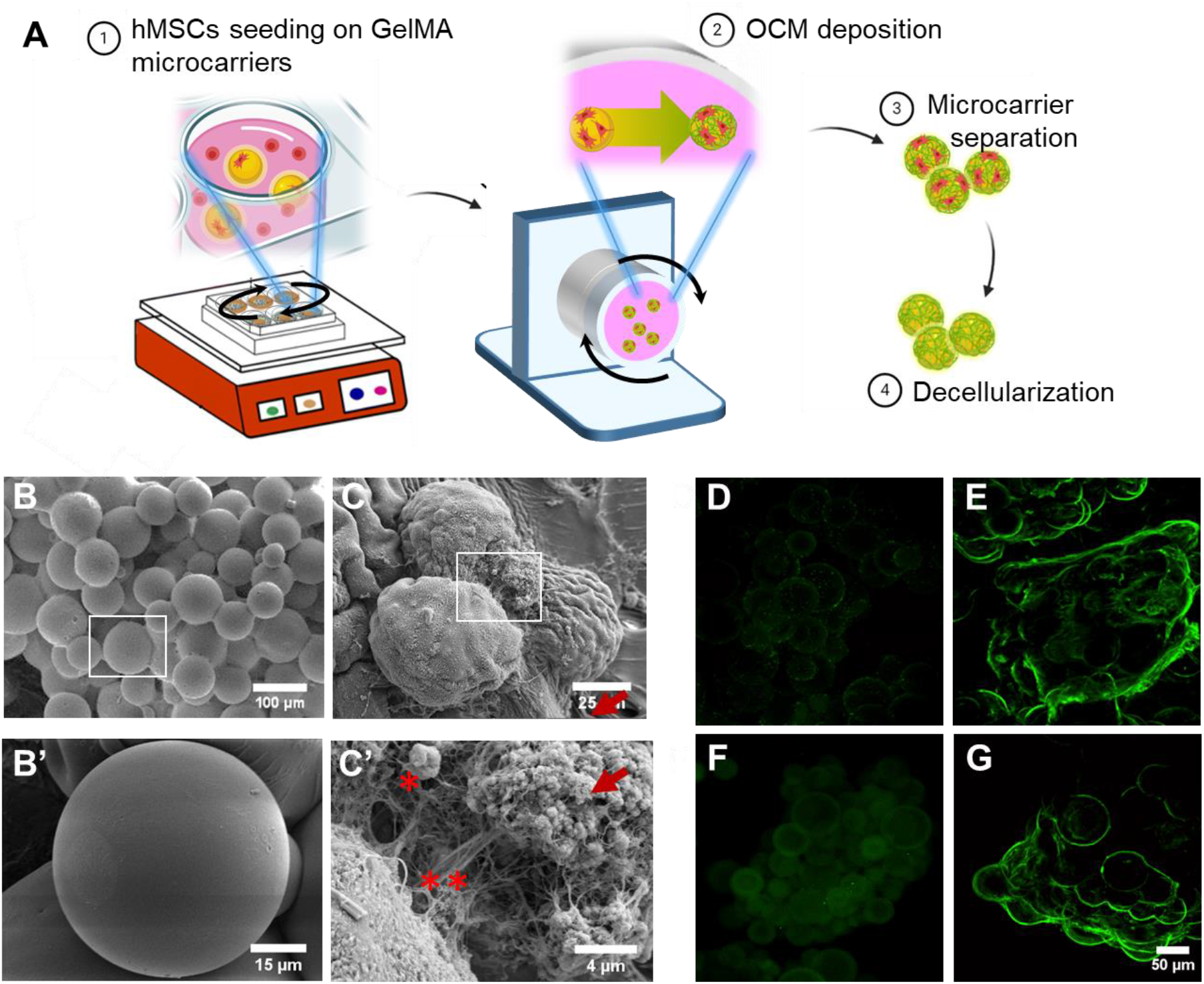
Preparation and characterization of decellularized OCM-GelMA microcarriers. A) Schematic diagram of the steps to obtaining OCM-coated microcarriers. B-C: Representative low- and high-resolution SEM micrographs of GelMA (B, B’) and OCM-GelMA (C, C’) microcarriers. D-G) Representative immunofluorescence images of GelMA (D,F) and OCM-GelMA (E,G) microcarriers with antibodies against collagen VI (D, E) and XII (F, G).

### OCM accelerates hMSC expansion on OCM-coated microcarriers

After decellularization, the resulting OCM-GelMA microcarriers were seeded with fresh hMSCs to investigate the ability of the decellularized OCM to maintain cell viability and promote expansion in CCM. Live/dead assays indicated comparable levels of cell viability on GelMA and OCM-GelMA microcarriers (Figures 3A-C). Over a period of 8 days of suspension culture, hMSCs expanded rapidly on OCM-GelMA microcarriers, reaching a peak of approximately 9×10^4^ cells/cm^2^ in 5 days (Figure 3D). In comparison, hMSCs on GelMA microcarriers required 8 days to achieve comparable numbers of cells (Figure 3D).

**Figure 3.**
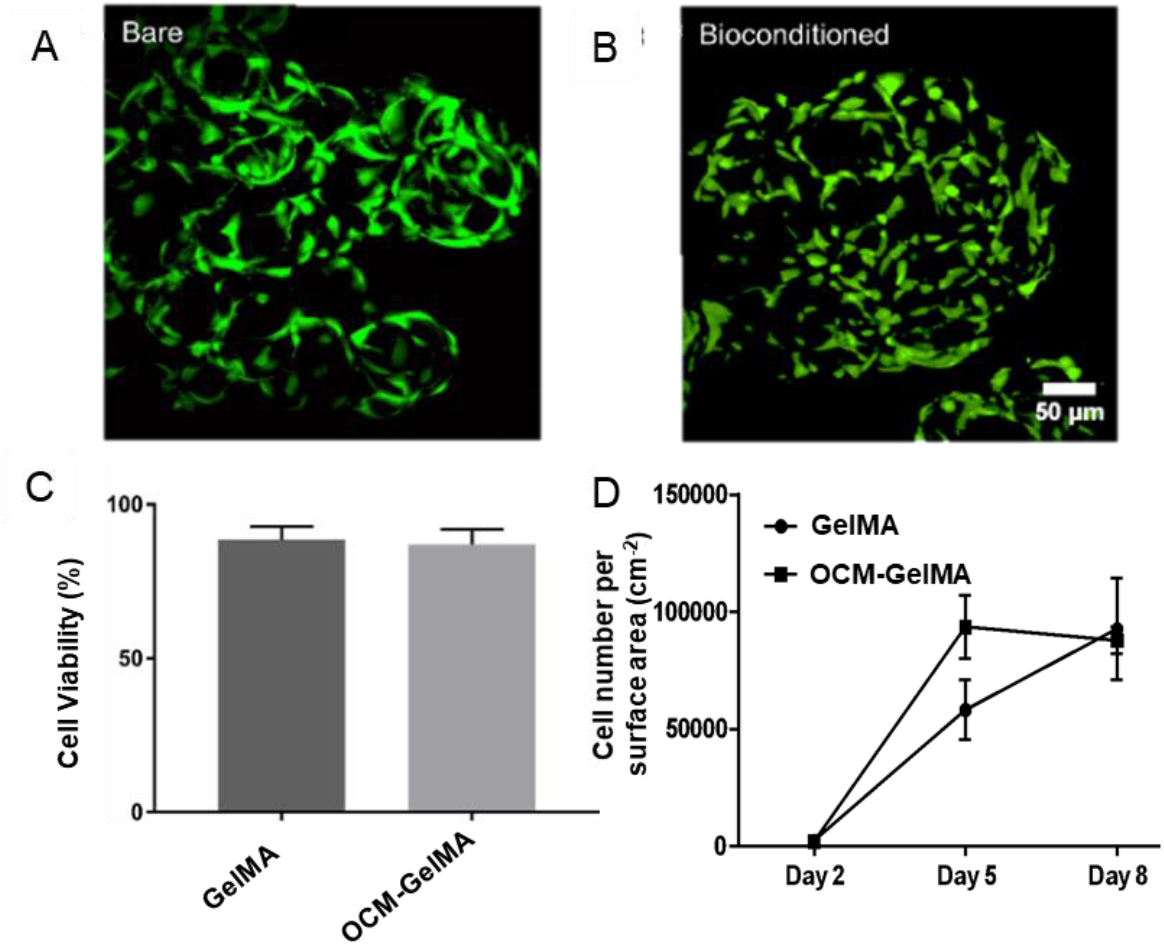
hMSC viability and expansion on microcarriers. A-C: Live/dead staining of hMSCs present on GelMA and OCM-GelMA microcarriers two days after seeding demonstrated an abundance of live cells and sparsely distributed dead cells. hMSCs were labeled with calcein-AM (green) to visualize live cells and propidium iodide (red) to visualize dead cells and seeded on top of bare (A) and OCM-coated (B) microcarriers, with the results summarized (C) (average ± s.d; N=3). **D:** cell numbers were determined using qRT-PCR quantification of GAPDH expression of hMSCs cultured on bare and OCM-coated microcarriers in the bioreactor over time (average ± s.d; N=3).

### Effects of OCM on the hMSC secretome

The regenerative potency of hMSCs has been associated with the secretion of several bioactive factors, including basic fibroblast growth factor (FGF-2), vascular endothelial growth factor (VEGF), transforming growth factor *β*1 (TGF-*β*1), and platelet derived growth factor-AA (PDGF-AA) [24, 25]. We compared the levels of secretion of multiple cytokines associated with osteogenesis and angiogenesis from hMSCs cultured on GelMA and OCM-GelMA microcarriers in suspension culture in the presence of GW9662, with hMSCs cultured on GelMA microcarriers in the absence of GW9662 serving as a negative control (Figure 4). OCM enhanced secretion of each of the cytokines tested, with significant differences detected for FGF-2, VEGF and TGF-*β*1, but not PDGF-AA. GW9662 treatment enhanced secretion of FGF-2 and the early osteogenic biomarker osteoprotegrin (OPG) for hMSCs on bare microcarriers, but the effect was only significant for FGF-2 secretion. Secretion of Dickkopf-1 (Dkk-1), an inhibitor of canonical Wnt signaling during terminal differentiation of osteoprogenitor cells [26, 27], was significantly reduced when hMSCs were cultured on OCM even though GW9662 treatment alone enhanced Dkk-1 secretion from hMSCs on bare microcarriers.

**Figure 4.**
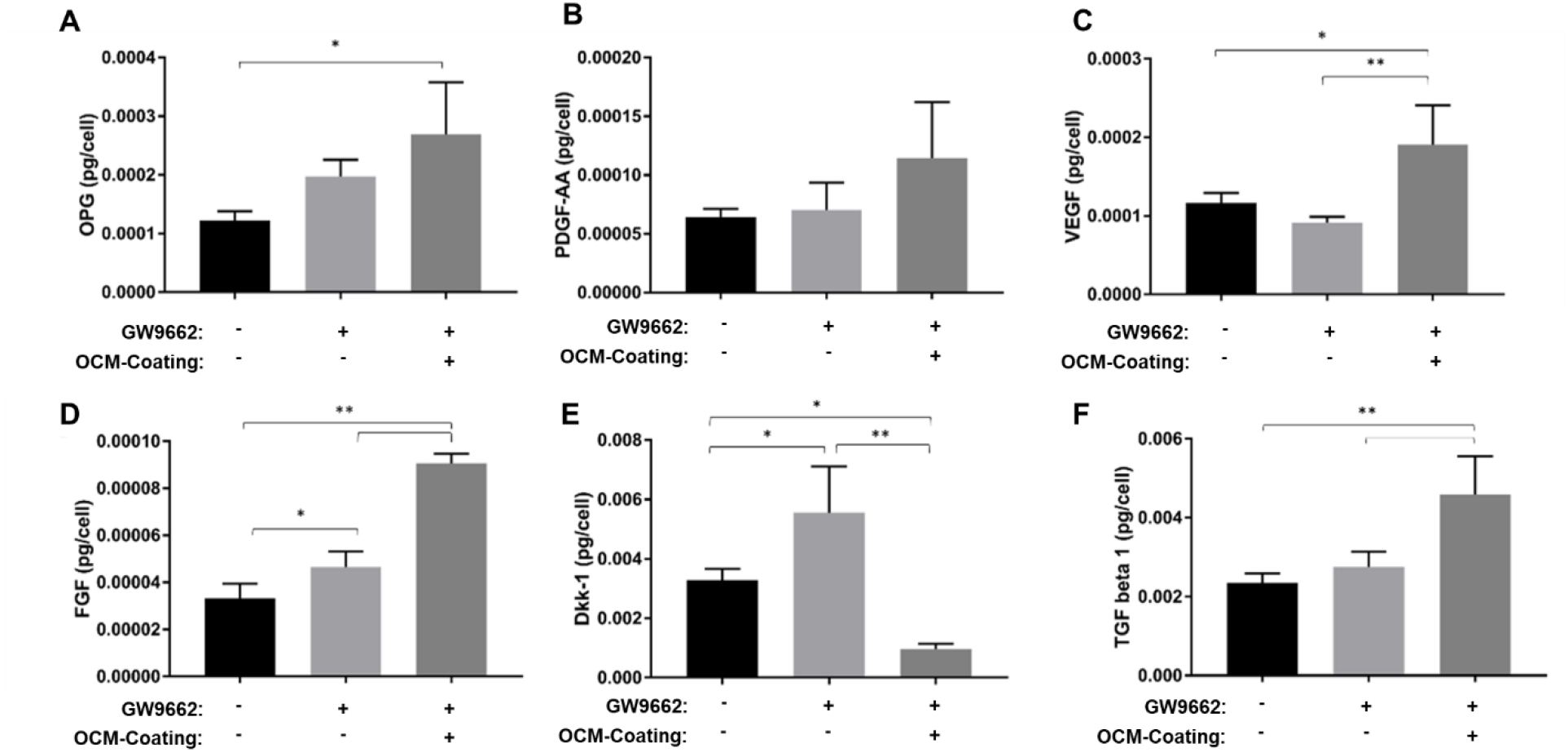
OCM-GelMA enhances osteogenic and angiogenic growth factor secretion. OPG (A), PDGF-AA (B), VEGF (C), FGF (D), Dkk-1 (E), and TGF beta 1 (F) secretion was measured from the medium in RWVs by ELISA on day 4. Values were normalized to cell number. Statistics are (n=3) one sided ANOVA and Tukey post test ^*, **^represents p<0.05, 0.01. All data presented as average ± s.d. All experiments performed in triplicate.

### Effects of OCM on the transcription of BMP-2 and Collagens VI and XII

We have previously demonstrated that OCM-coated gelatin foam scaffolds stimulate the expression of collagens type VI and XII, as well as bone morphogenetic protein 2 (BMP-2), in hMSCs treated with GW9662 [10]. In suspension culture, hMSCs grown on GelMA or OCM-GelMA microcarriers showed a marginal upregulation of collagen type VI gene expression when treated with GW9662, though the results were not statistically significant (Figure 5A). Administration of GW9662 to hMSCs on GelMA microcarriers resulted in an increase in collagen XII expression (Figure 5B). In contrast, the expression of collagen XII was suppressed in hMSCs cultured on OCM-GelMA microcarriers. The expression of BMP-2 was highly upregulated in hMSCs cultured on OCM-coated microcarriers (Figure 5C).

**Figure 5.**
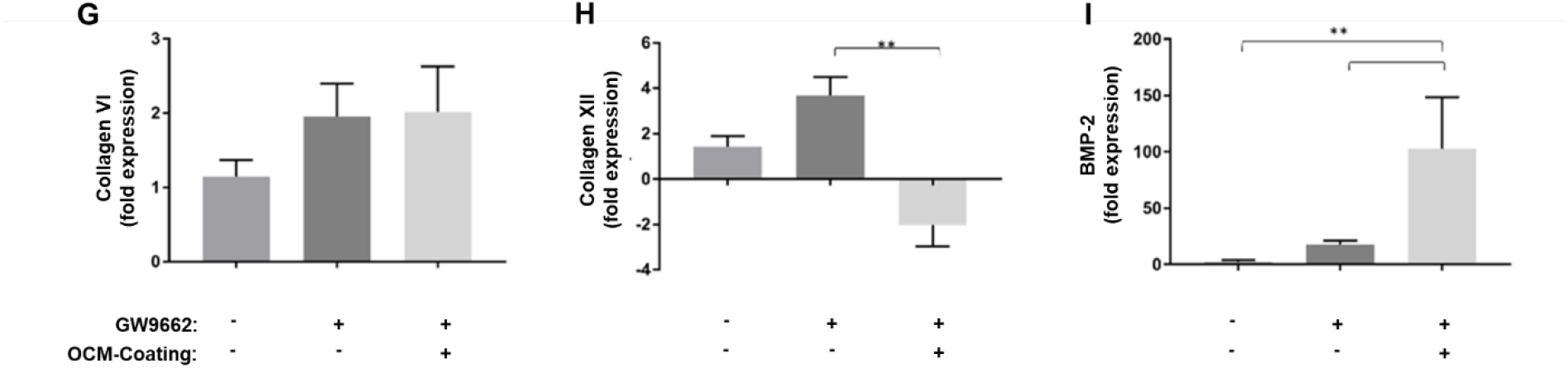
Effects of OCM and GW9662 on collagen and BMP-2 gene expression. Quantitative PCR was used to quantify the effects 10uM GW9662 and OCM on transcription of collagen VI (A) and collagen XII (B) on Day 8. (C) GW9662 and OCM-GelMA synergistically upregulated BMP-2 expression on Day 8. Data is presented as fold-change in gene expression relative to hMSC monolayers in Complete Culture Media. Statistics are (n=3) one sided ANOVA and Tukey post test ^**^represents p<0.005. All data presented as average ± s.d. All experiments performed in triplicate.

### Bone regeneration in murine calvarial defect model

We previously demonstrated that the bone regenerative capabilities of OEhMSCs is substantially augmented by co-administration of OCM using a murine calvarial defect model [6, 7]. In those studies, OCM was harvested from OEhMSC monolayers. In the present study, GelMA and OCM-GelMA microcarriers were delivered, with or without OEhMSCs, into circular full-thickness calvarial defects and imaged by digital radiography 4 weeks post-surgery (Figure 6A). The extent of healing induced when OCM-GelMA microcarriers were administered alone was moderately higher than that induced by GelMA microcarriers and mock negative controls that did not receive any materials (Figure 6B). Co-administration of OEhMSCs with GelMA or OCM-GelMA microcarriers resulted in statistically increased healing compared to mock controls. Co-administration of OEhMSCs with OCM-GelMA microcarriers resulted in healing that was significantly higher than co-administration with GelMA microcarriers. As a positive control, BMP-2 was delivered in a gelatin sponge and this resulted in comparable healing to that induced by OEhMSCs administered with OCM-GelMA microcarriers.

**Figure 6.**
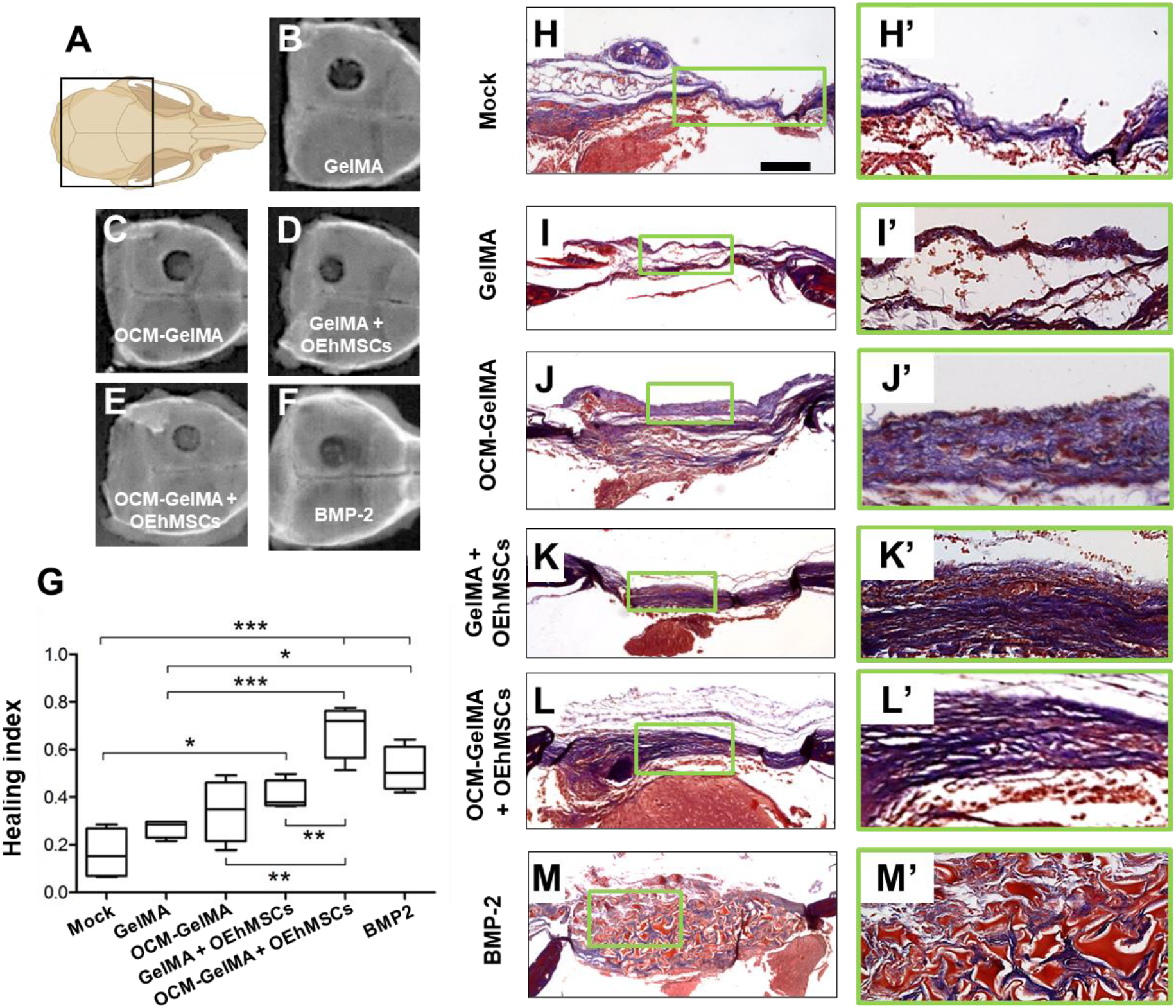
Bone healing in a murine calvarial lesion model. Schematic (A) and representative digital radiographs after 4 weeks of healing (B-F) of mock surgery (negative control) and implantations of GelMA or OCM-GelMA microcarriers with or without OEhMSCs. The positive control consisting of 50 µg BMP-2 on a gelatin sponge (BMP-2) was administered without OEhMSCs. (G) Summary of RHI values at each condition where a value of 0 represents no healing and 1 represents healing equivalent to the contralateral side. Low (H-M) and medium (H’-M’) power micrographs of sagittal sections of calvaria at defect midpoint for each condition. Scale bar in panels H-M is 1 mm. Paraffin sections were stained with Masson’s Trichrome.Statistics: n=4, analyzed by ANOVA and Tukey post-test: ^*^p<0.05, ^**^p<0.01, ^***^p<0.005.

Histological staining, performed using the same tissue that was imaged by radiography, provided additional insight into the bone healing induced by the various conditions tested (Figure 6H-M). Mock defects (Figure 6H) resulted in a thin fibrotic scar with very little new tissue. Implantation of GelMA or OCM-GelMA microcarriers alone (Figures 6I vs 6J) stimulated the generation of thicker fibrous tissue, but bone formation was not evident. GelMA microcarriers administered with OEhMSCs induced formation of bone (Figure 6K) combine with fibrous tissue (Figure 6K’). OCM-GelMA microcarriers administered with OEhMSCs generated significantly more new bone as indicated by the dark blue staining (Figure 6L) with less indiction of fibrous tissue (Figure 6L’). BMP-2 induced the formation of numerous diffusely distributed isolates of immature bone tissue in between patches of gelatin foam (Figure 6M).

### Bone regeneration in murine femoral defect model

To assess endochondral bone repair, we employed a pin-stabilized murine femoral defect model [10]. Qualitative observation of µCT scans indicated minimal signs of healing after 5 weeks of healing in cases where GelMA microcarriers were administered with or without OEhMSCs (Figure 7B and D). A small amount of bone formation was observed when OCM-GelMA microcarriers were administered without OEhMSCs (Figure 7C). In contrast, significant bone formation was observed with femora that received OCM-GelMA microcarriers with OEhMSCs (Figure 7E). Unexpectedly, extensive bone growth could be observed not only within the defect, but also throughout the femora at distal sites, suggesting that the OCM-GelMA microcarriers and OEhMSCs had distributed themselves along the axis of the bone and their osteoanabolic activity was not confined to the defect (Figure 7E). Therefore, the volume of the entire femur was measured for each sample to quantify the overall impact on bone formation. Bone surface-to-volume ratio is an indicator of *de novo* bone compactness and can be extrapolated as an indicator of bone strength. These data confirmed that OCM-GelMA microcarriers are advantageous for stimulation of bone formation in murine femora.

**Figure 7.**
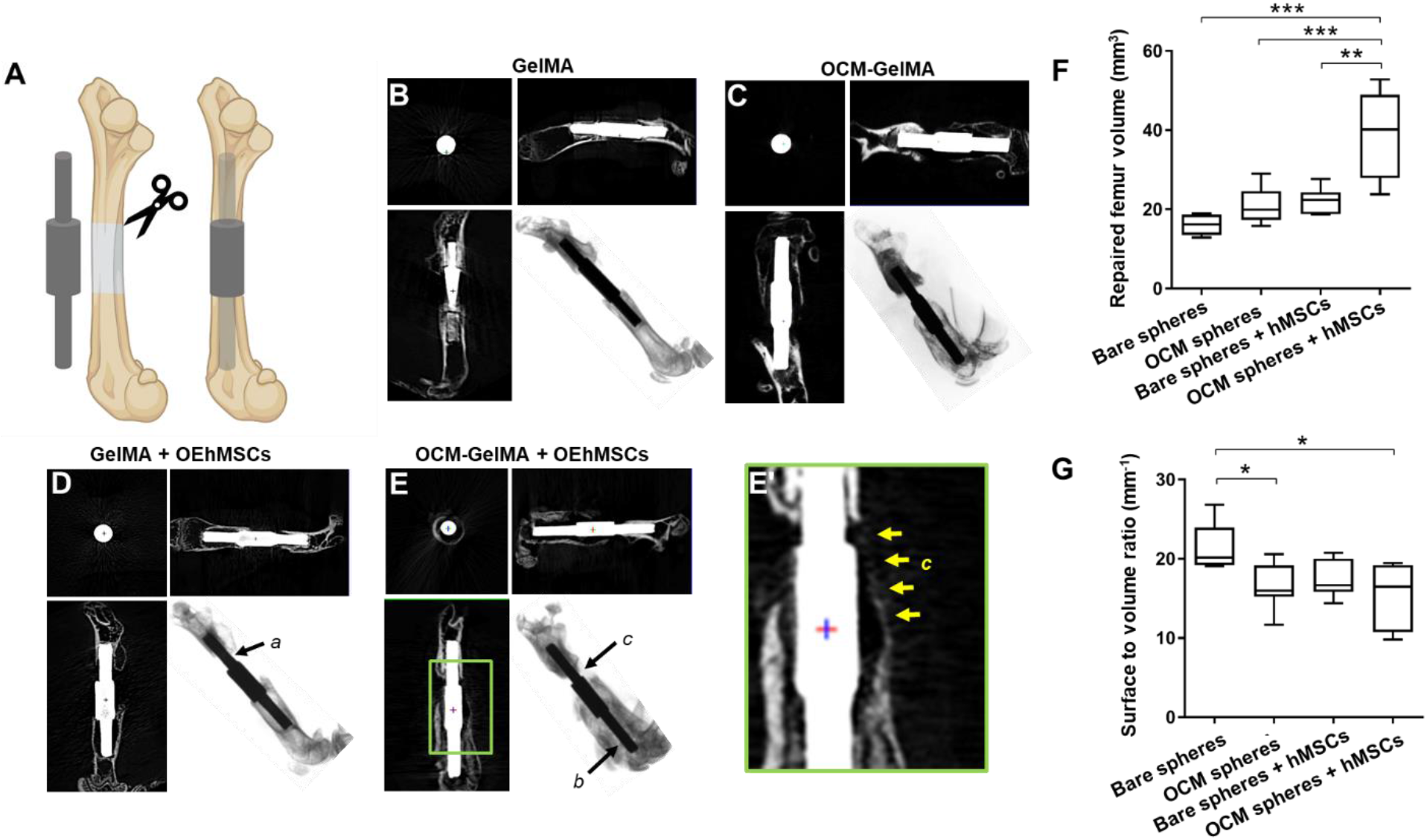
Stimulation of bone formation by OCM-GelMA microcarriers and OEhMSCs in a murine critical femoral defect model. (A) Schematic of the pin-stabilized segmental femoral defect. (B-E) Micro-CT-generated orthogonal images and x-ray scans. Arrow a indicates a poor healing response with a deficit of bone at the collar of the pin, arrow b indicates extensive bone growth at a distance from the defect, and arrow c indicates bridging over the entire defect with new bone in contact with the margin of the defect. (F) Whole femur volume (G) and surface-to-volume ratio as measured by µCT. (E’) Close-up view from (E) showing partial bridging (arrowed). Statistics: n=6-7, analyzed by ANOVA with Tukey post-test. ^*^=p<0.05, ^**^=p<0.01, ^***^=p<0.005.

Quantitative measures of repaired volume derived from these images showed significantly higher healing in femoral defects with OEhMSCs administered with OCM-GelMA microcarriers than any of the other conditions tested (Figure 7F). The defects treated with OCM-GelMA microcarriers administered with or without OEhMSC exhibited greater bone compactness compared to defects treated with GelMA microcarriers alone (Figure 7G)

Histological staining of the tissues imaged in Figure 8 indicated that GelMA microcarriers had a negligible effect on bone healing (Figure 8A), while OCM-GelMA microcarriers appeared to induce the formation of a thin layer of compact, cortical-like bone tissue, as indicated by the dense organization and dark blue staining, with no sign of trabecular structures and minimal cellularity (Figure 8B, E and F). GelMA microcarriers administered with OEhMSCs formed tissue with a high degree of cellularity (Figure 8I), fibrotic deposits and diffuse signs of bone formation (Figure 8C, G and H). The femoral defects that received OCM-GelMA microcarriers and OEhMSCs showed an increase in bone density around the collar indicating bone regeneration as indicated by the dark blue staining (Figure 8D). These samples displayed a spongy appearance, suggesting the formation of a trabecular network (Figure 8 D, J and K) and signs of both osteoblast (Figure 8L, *arrows*) and osteoclast (Figure 8L, *asterisks*) activity.

**Figure 8.**
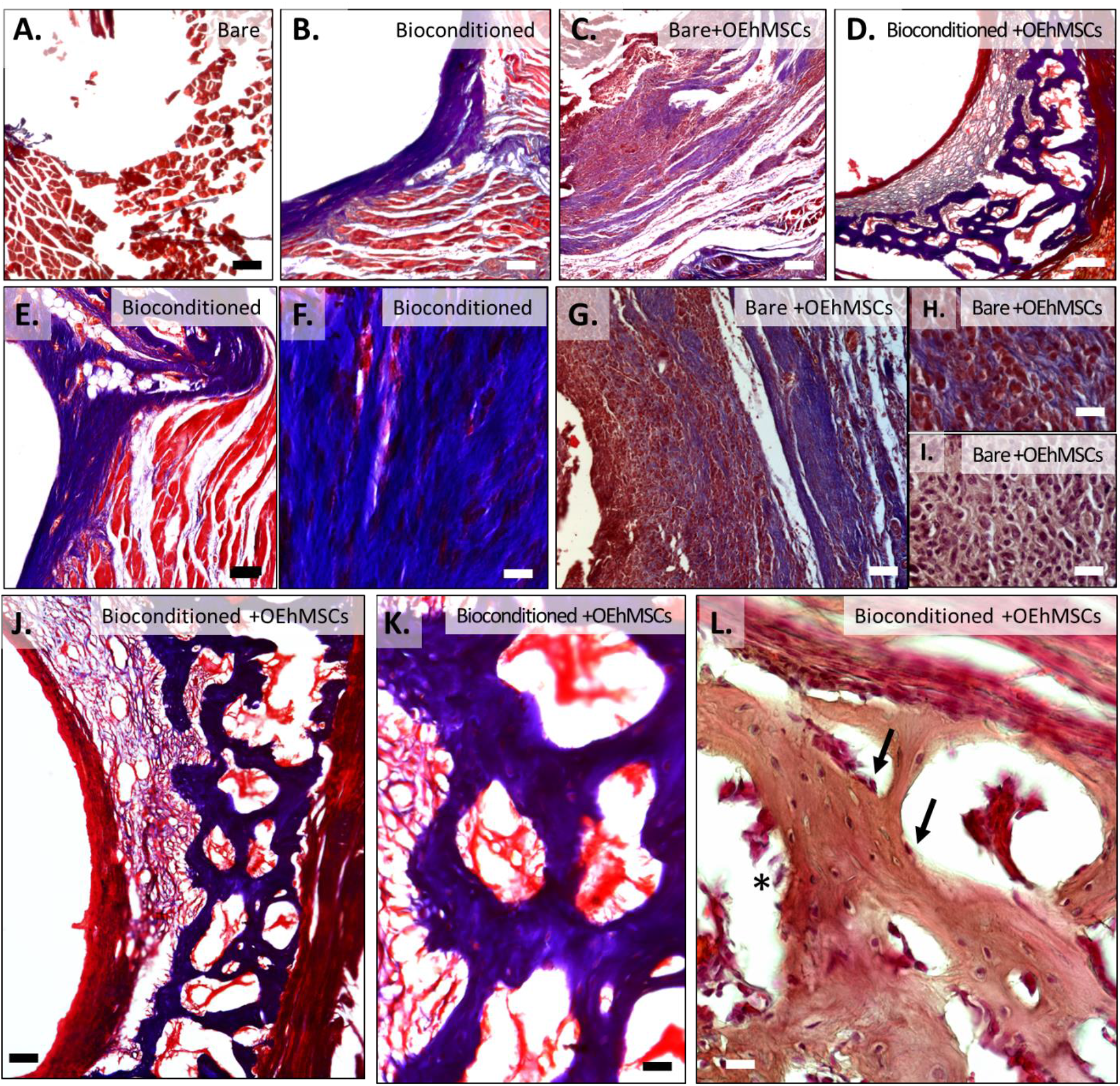
Histology of axial sections of femurs at defect midpoints treated with microcarriers and OEhMSCs. (A-D) Low power micrograph (4x, bar=250 µm). (E, G, J) Mid-power micrographs (10x, bar=100 µm). (F, H, I, K, L) High-power micrographs (20x, bar=25 µm). (A) GelMA microcarriers, (B, E, F) OCM-GelMA microcarriers, (C, G, H, I) GelMA microcarriers with OEhMSCs, (D, J, K, L) OCM-GelMA microcarriers with OEhMSCs. I and L stained with hematoxylin and eosin, all others stained with Masson’s trichrome. Arrows indicate osteoblasts and asterisks indicate patches of osteoclasts.

## DISCUSSION

Cell therapy strategies using bone marrow cells represent an attractive alternative to autologous bone grafting for treating significant critical size osseous defects when bone healing ability is limited, such as in elderly patients or diseases which have a high recurrence even after surgical intervention such as aneurymal bone cysts and congenital pseudarthrosis [28]. Defects in long bones and craniomaxillofacial bones heal by different mechanisms with cells derived from different developmental lineages, leading to the argument that the success of therapeutic approaches may differ depending on which type of bone is being treated [29]. This knowledge motivated assessment of OCM-GelMA microcarriers in both femoral and calvarial critical defect models. The OCM-GelMA microcarriers described in this study represent an osteogenic vehicle for delivering hMSCs into osseous defects to accelerate bone healing to an extent at least comparable to BMP-2, a potent bone graft material used in high-risk bone grafting cases. Our results demonstrate that OCM-GelMA microcarriers induce a pro-regenerative phenotype in hMSCs treated with GW9662, characterized by increased secretion of OPG, VEGF, FGF-2 and TGF-β1 and increased expression of BMP-2. Further, implantation of OCM-GelMA with OEhMSCs resulted in greater bone deposition in both long bone and calvarial critical size defects, providing evidence that OCM-GelMA microcarriers have potential as an osteogenic vehicle for percutaneous delivery of bone marrow cells for accelerating bone healing in patients.

The paracrine effects of hMSCs are well established [30], which has motivated clinical investigations of incorporating conditioned media from hMSCs into bone grafts [31].

These investigators previously demonstrated that conditioned media was effective in accelerating calvarial healing in a rat model and detected significant amounts of insulin-like growth factor (IGF)-1 and VEGF, but not FGF-2, PDGF-BB or BMP-2 in the conditioned media. In the presence of GW9662, we detected VEGF, PDGF-AA, FGF-2, and TGF-β in the conditioned media of hMSCs grown on GelMA-OCM microcarriers.

Further, the concentrations of VEGF, FGF-2, and TGF-β are significantly higher than when hMSCs were grown on GelMA microcarriers. These results suggest that the bone healing potency of hMSCs administered with GelMA-OCM is at least in part due to the secretion of these paracrine factors. We previously demonstrated that OCM delivered directly into calvarial defects extended the duration of retention of implanted OEhMSCs [7]. Since the implanted OEhMSCs were confined to OCM rather than the surface of the remodeling bone, it appears that paracrine effects of the OEhMSCs are more relevant than direct contribution to tissue repair [7].

The results of this study suggest that the composition of extracellular matrix proteins on the surface of microcarriers dictate the behavior of attached hMSCs. In addition to influencing the secretion of osteogenic and angiogenic paracrine factors, culturing GW9662-treated hMSCs on OCM-GelMA microcarriers resulted in no increase in collagen VI expression and a decrease in collagen XII expression. One potential explanation for this result is that the presence of collagens VI and XII suppresses further expression of these collagen isoforms since they are already present. On the other hand, expression of BMP-2 is dramatically upregulated on OCM-GelMA microcarriers relative to bare GelMA microcarriers, suggesting that the protein synthesis is redirected to production of a protein directly associated with bone formation. This is consistent with increased bone repair induced by GelMA-OCM vs bare GelMA microcarriers. Thus collagens VI and XII may stimulate more efficient bone healing by hMSCs.

In one study involving patients with distal tibial fractures treated with intramedullary nail, percutaneous delivery of isolated hMSC in combination with demineralized bone matrix (DBM) and platelet-rich plasma was reported to accelerate fusion rates and reduce delayed union relative to control cases lacking the bone graft [32]. DBM is reported to contain BMPs responsible for osteoinductive properties, though variability in growth factor concentration between products and between lots of a single product have limited its widespread use for nonunion treatment [33]. Based on the results presented herein, OCM-GelMA microcarriers may represent an improvement over DBM in terms of composition and osteogenic activity. OCM-GelMA microcarriers are coated with collagen types VI and XII, which we have shown to be critical to the potency of OCM in healing calvarial [34] and femoral [10] defects. The mechanisms by which collagens VI and XII contribute to bone regeneration remain unclear, but their prevalence in developing and regenerating bone relative to homeostatic adult bone [9, 35-37] and contributions to bone malformations when absent strongly implicate roles for these collagen isoforms in bone regeneration.

Delivery of bone marrow cells and BMP-2 have each been extensively investigated for use in craniofacial reconstruction [38]. BMP-2 delivered in a gelatin sponge has been used successfully in the clinic for bone grafting of the alveolar ridge in cases with high risk of poor bone healing outcomes [39]. We observed that BMP-2 induced comparable levels of calvarial healing as GelMA-OCM delivered with OEhMSCs, but the bone associated with BMP-2 was relatively less compact. This result is consistent with our recent report comparing calvarial healing associated with BMP-2 versus OCM harvested from induced pluripotent stem cell-derived MSCs in conventional monolayer culture [34].

Microcarrier culture represents a promising strategy for producing large quantities of hMSCs for clinical trials. Light polycaprolactone microcarriers have been used to expand hMSCs in stirred cultures [40]. In the study, hMSCs on the microcarriers produced higher levels of multiple cytokines and exhibited greater osteogenic differentiation than hMSCs in monolayer culture. Further, the hMSCs on the microcarriers induced greater healing in a rat calvarial defect than autograft 16 weeks post-surgery. A potential advantage to using GelMA microcarriers is that GelMA can be degraded by proteolytic enzymes relatively quickly to facilitate replacement by compact bone, while polycaprolactone degrades far more slowly.

Agitation of emulsions in a rapidly stirred bath represents a straightforward way to produce microdroplets for use in bone regeneration. Annamalai et al. [41] encapsulated murine adipose MSCs in a mixture of chitosan, hydroxyapatite microparticles and collagen Type I emulsified in a stirred polydimethylsiloxane bath to result in microtissues of comparable size to those used in the present study. Pre-osteogenic differentiation of the murine adipose MSCs dramatically enhanced calvarial healing after 12 weeks, which is consistent with our previous study showing that OEhMSCs significantly improved healing of calvarial defects relative to undifferentiated hMSCs [6]. Our experiments quantified the extent of calvarial healing at 4 weeks, making direct comparisons to those reported by Annamalai et al. [41] are unclear. Instead, we demonstrated a comparable extent of healing, but with more compact bone, relative to that induced by BMP-2, a clinically relevant control.

Our flow-focusing method used to generate microcarriers generation has both merits and limitations. Flow-focusing emulsification can greatly reduce variance in droplet size compared to stirred-bath emulsification [42-44]. Stereolithographic 3D printing facilitates reproducible fabrication of the coaxial nozzle. In this system, droplet pinch-off is driven by shear forces from the outer oil flow. Consequently, droplet size can be controlled by modulating the ratio of oil-to-aqueous flow rates. Any variation in this flow rate ratio will contribute to variation in the droplet sizes, however. Step emulsification has emerged as a more practical method for droplet formation that is driven by flow instability with the droplet size now dependent on the orifice geometry rather than shearing forces [45]. We recently reported the use of step emulsion with 100 parallel orifices for high-throughput generation of GelMA microcarriers with smaller variance in size than in the current study [46]. Due to the independence on oil-to-aqueous flow rate ratio, 3.3-fold greater microcarrier production rate was achievable with an oil-to-aqueous flow ratio of only 2 [46].

## AUTHOR CONTRIBUTIONS

C.H.: Conceptualization, Methodology, Formal analysis, Investigation, Writing (Original Draft, Review and Editing), Visualization; S.J.: Conceptualization; Methodology; Formal analysis; Investigation; E. Mondragon, S.P., E.McNeill, C.C., D.T., B.C., N.S., A.D.: Investigation, Writing (Review); J.K.: Conceptualization, Writing (Review), Project administration, Funding acquisition; C.A.G. and R.K.: Conceptualization, Methodology, Formal analysis, Writing (Review and Editing), Visualization, Project administration, Funding acquisition.

